## Supplementary material for "Phospho-islands and the evolution of phosphorylated amino acids in mammals": File containing all supplementary figures

### Supplementary materials for the article "Phospho-islands and the evolution of phosphorylated amino acids in mammals"

#### 1. Supplementary Figures

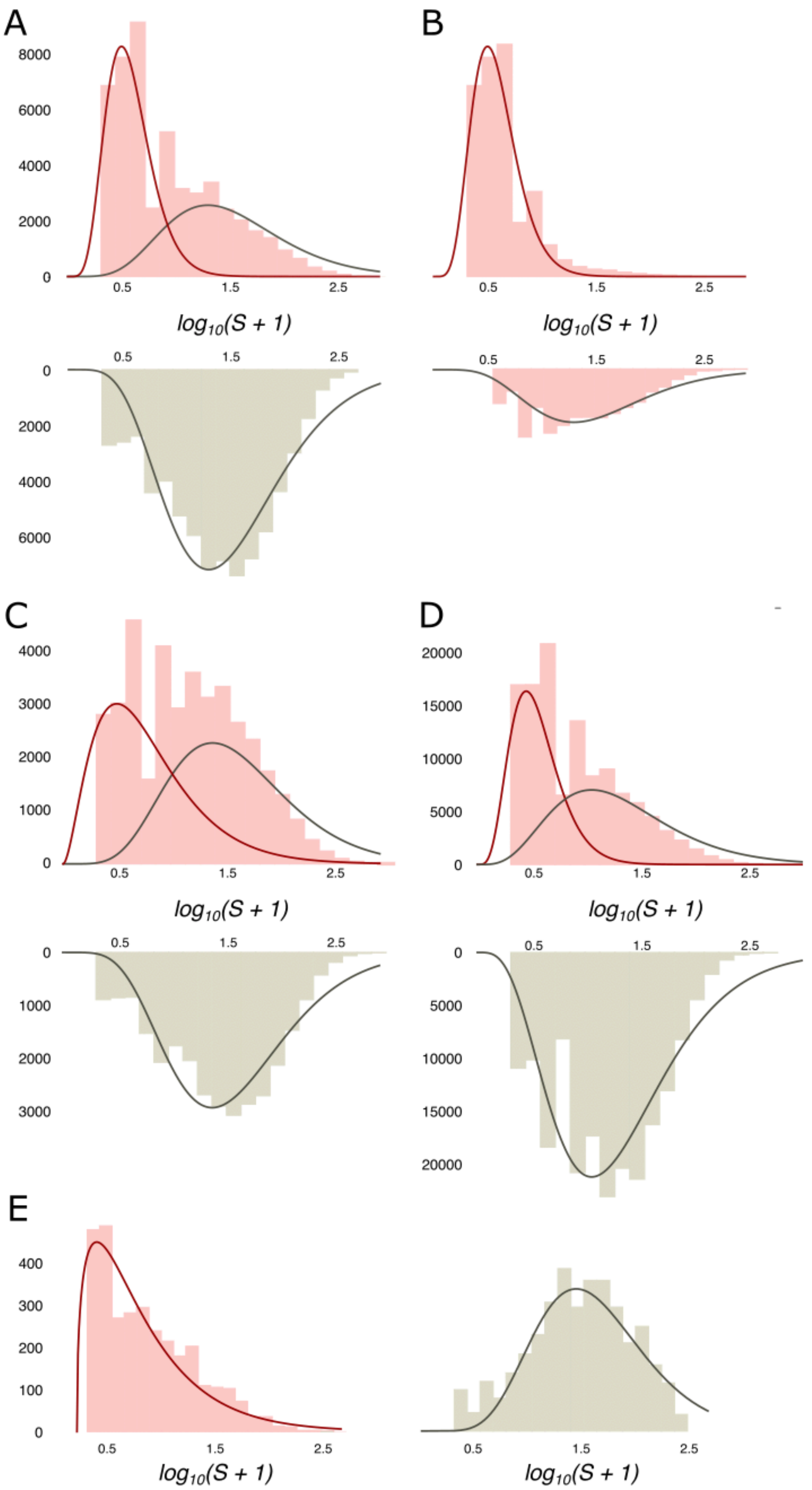


**Supplementary Figure S1 | Phospho-island analysis in various datasets. (A, B)** Prediction of phospho-islands for the mouse dataset. **(A)** (Above) The distribution of $\log_{10}(S + 1)$ values (pink histogram; see the main text for definitions) and its decomposition in two gamma distributions, for phospho-islands (red curve) and for individual phosphosites (red curve). (Below) $\log_{10}(S + 1)$ values for non-phosphorylated STY amino acids randomly sampled from DRs with the same sample size and amino acid content as in the HMR dataset. **(B)** (Above) The distribution of $\log_{10}(S + 1)$ values for phosphosites predicted to be in phospho-islands. (Below) $\log_{10}(S + 1)$ values for predicted individual phosphosites. **(C)** Histograms of $\log_{10}(S + 1)$ values for real (pink) and randomly assigned (grey) phosphosites with respective gamma-approximations for the rarefied human dataset. **(D)** Histograms of $S$ values for real (pink) and randomly assigned (grey) phosphosites with respective gamma-approximations for the human dataset. **(E)** Distributions of the $\log_{10}(S + 1)$ values of phosphosites located in ORs (pink) and the distribution of $\log_{10}(S + 1)$ for STY amino acids randomly sampled from ORs (grey) with respective gamma-approximation. Note the good fit of a single gamma-distribution for the $\log_{10}(S + 1)$ values observed in ORs.

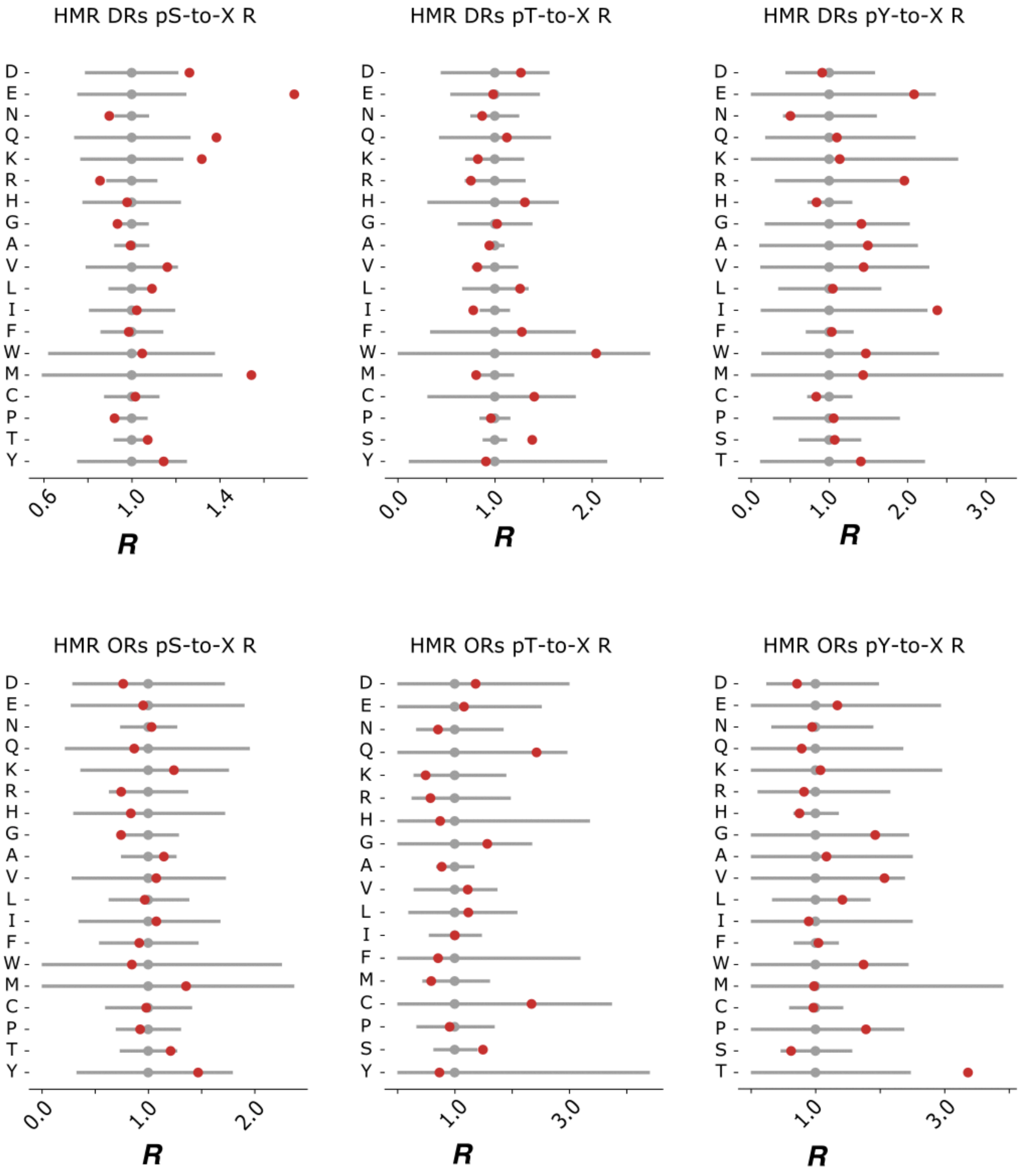

**Supplementary Figure S2 | Comparison of the mutational patterns observed in HMR phosphosites located in disordered and ordered regions.** *R* values are defined in the main text. 95% two-tail confidence intervals are generated from the chi-squared distribution with the Bonferroni correction. Red dots mark the actual *R* values. Grey dots represent the respective *R* values of non-phosphorylated STY amino acids.

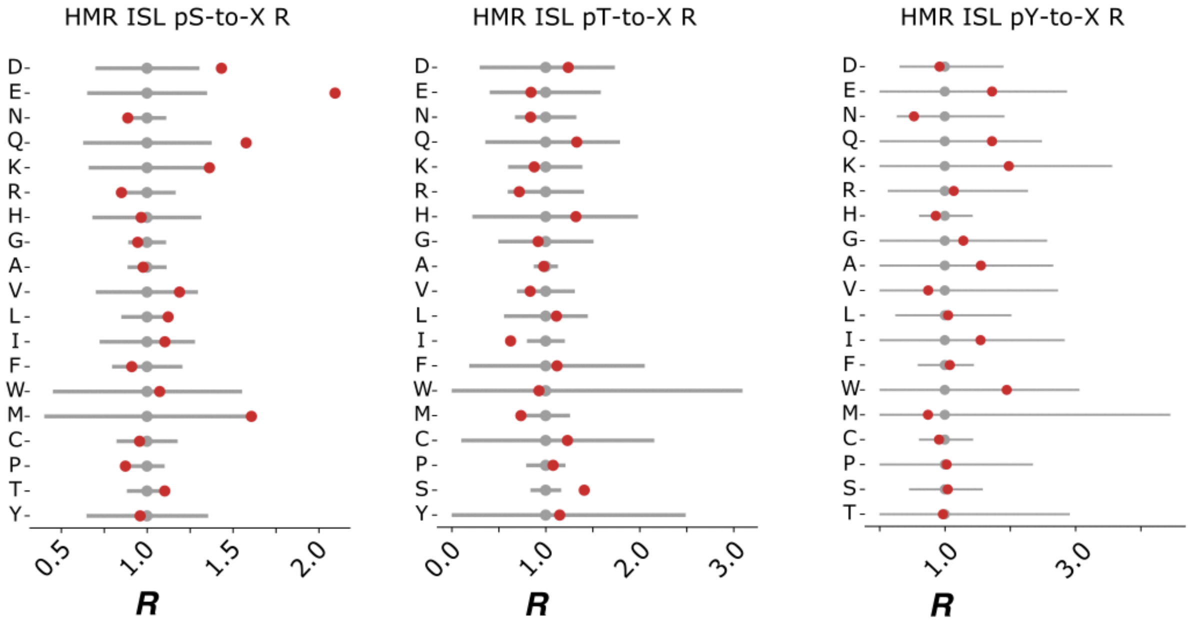


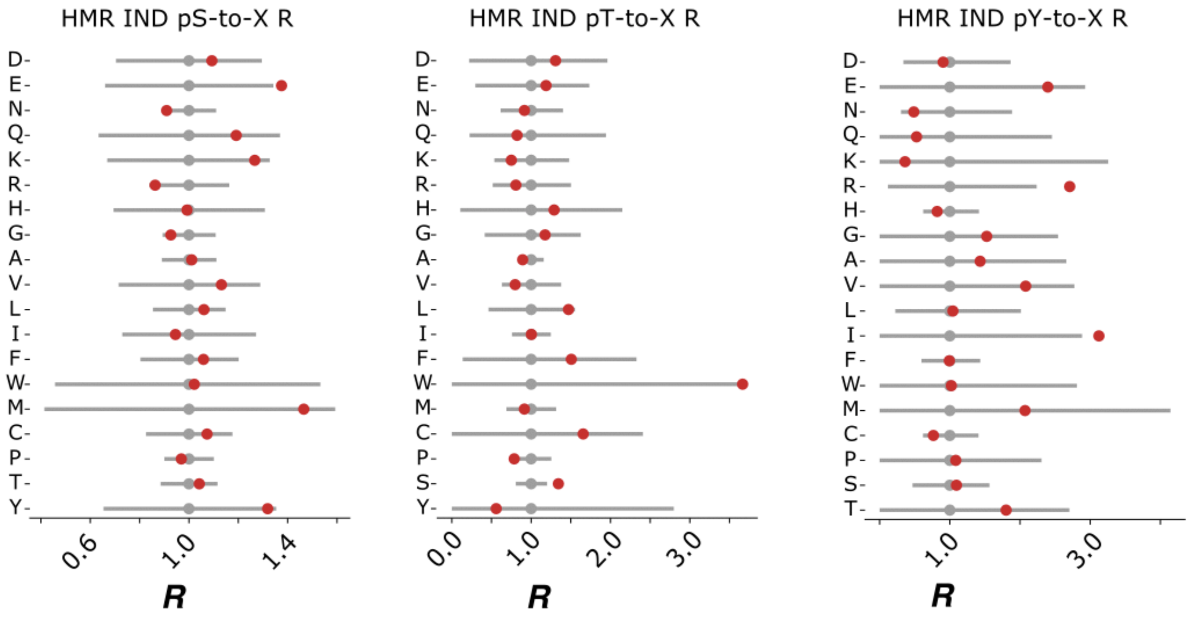


**Supplementary Figure S3 | Comparison of the mutational patterns observed in HMR phosphosites located in phospho-islands *vs.* individual ones.** Notation as in Suppl. Fig. S2.

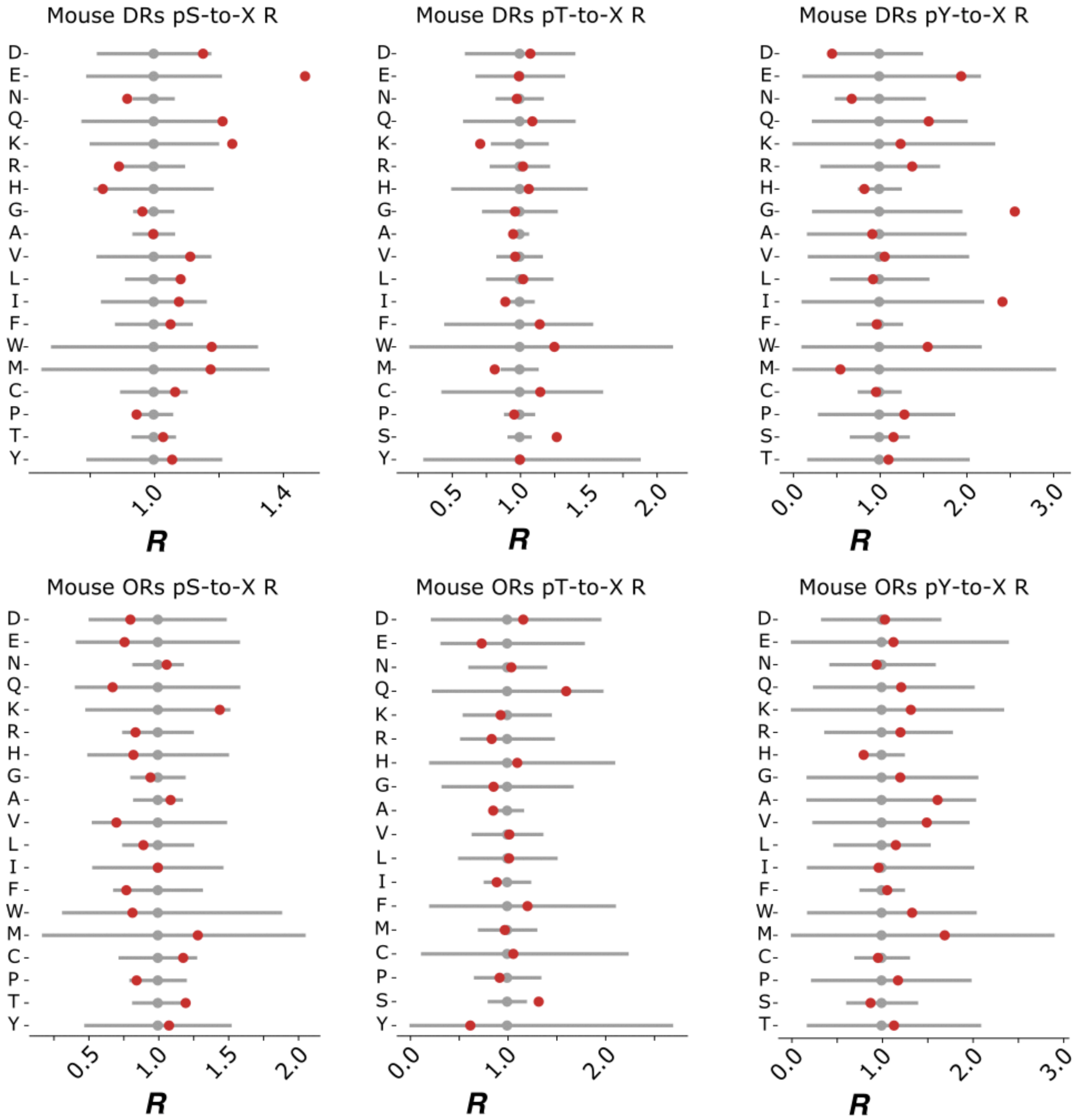


**Supplementary Figure S4 | Comparison of the mutational patterns observed in mouse phosphosites located in disordered and ordered regions.** Notation as in Suppl. Fig. S2.

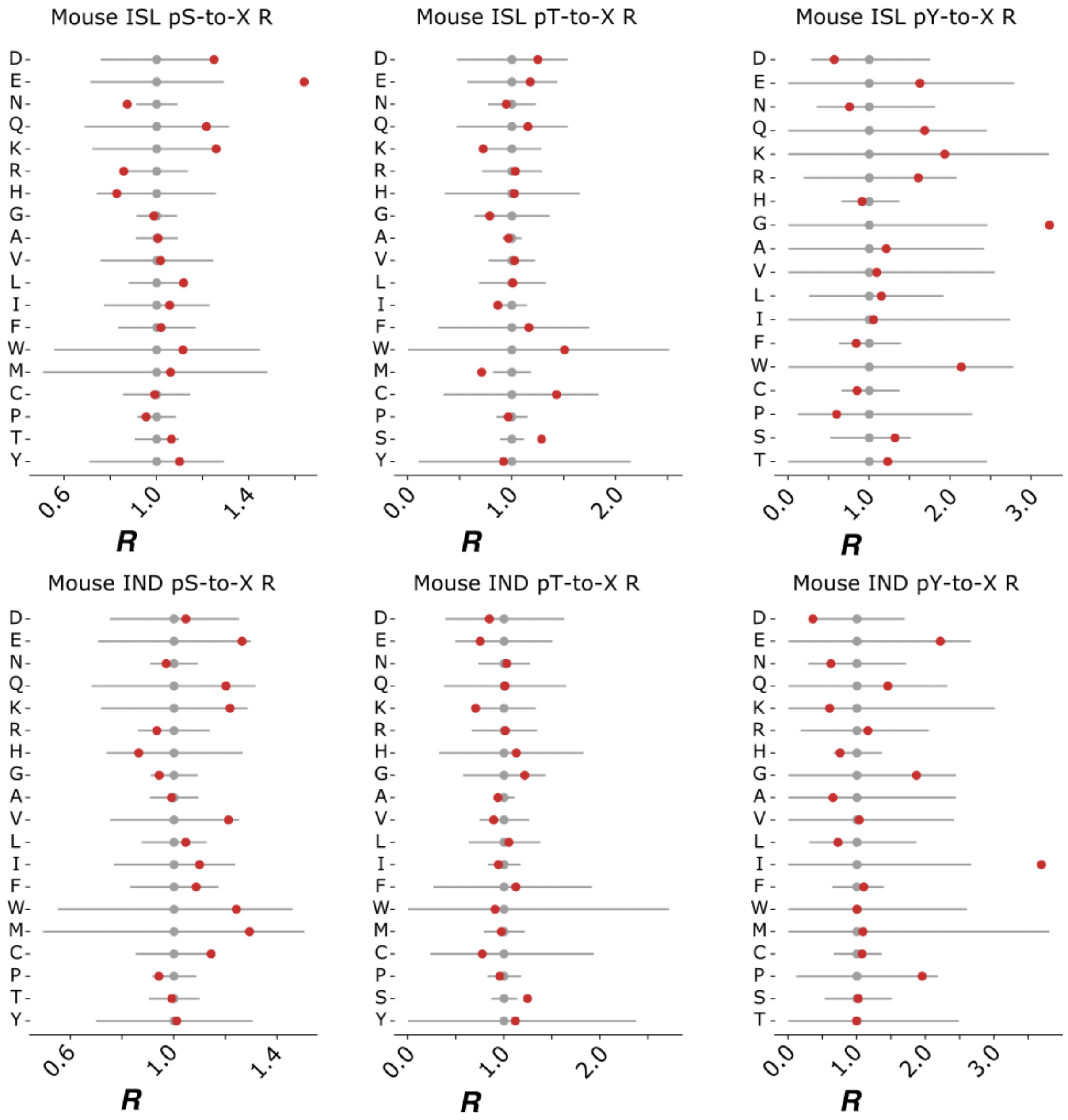


**Supplementary Figure S5 | Comparison of the mutational patterns observed in mouse phosphosites located in phospho-islands *vs.* individual ones.** Notation as in Suppl. Fig. S2.

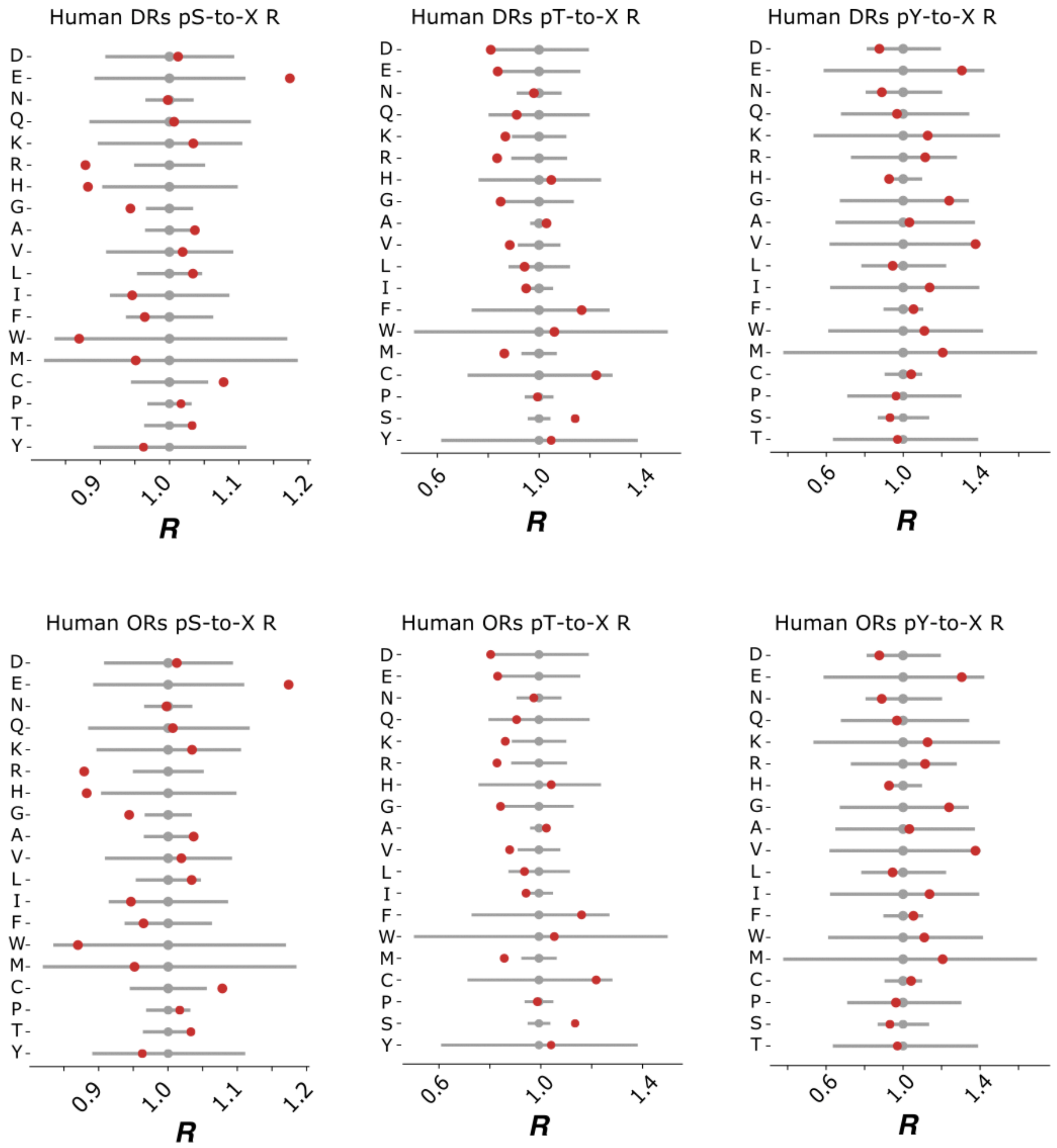


**Supplementary Figure S6 | Comparison of the mutational patterns observed in human phosphosites located in disordered and ordered regions.** Notation as in Suppl. Fig. S2.

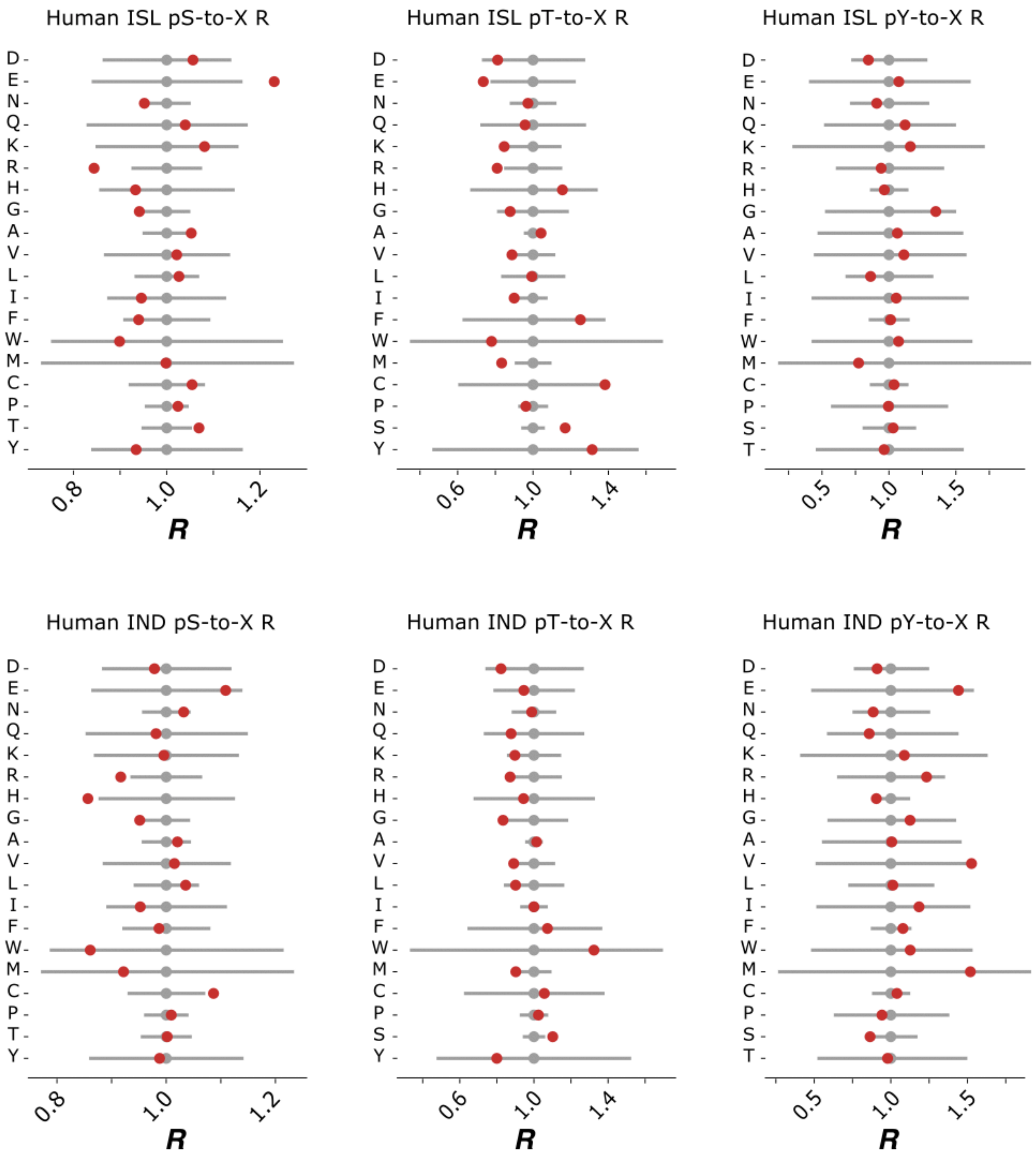


**Supplementary Figure S7 | Comparison of the mutational patterns observed in human phosphosites located in phospho-islands *vs.* individual ones.** Notation as in Suppl. Fig. S2.

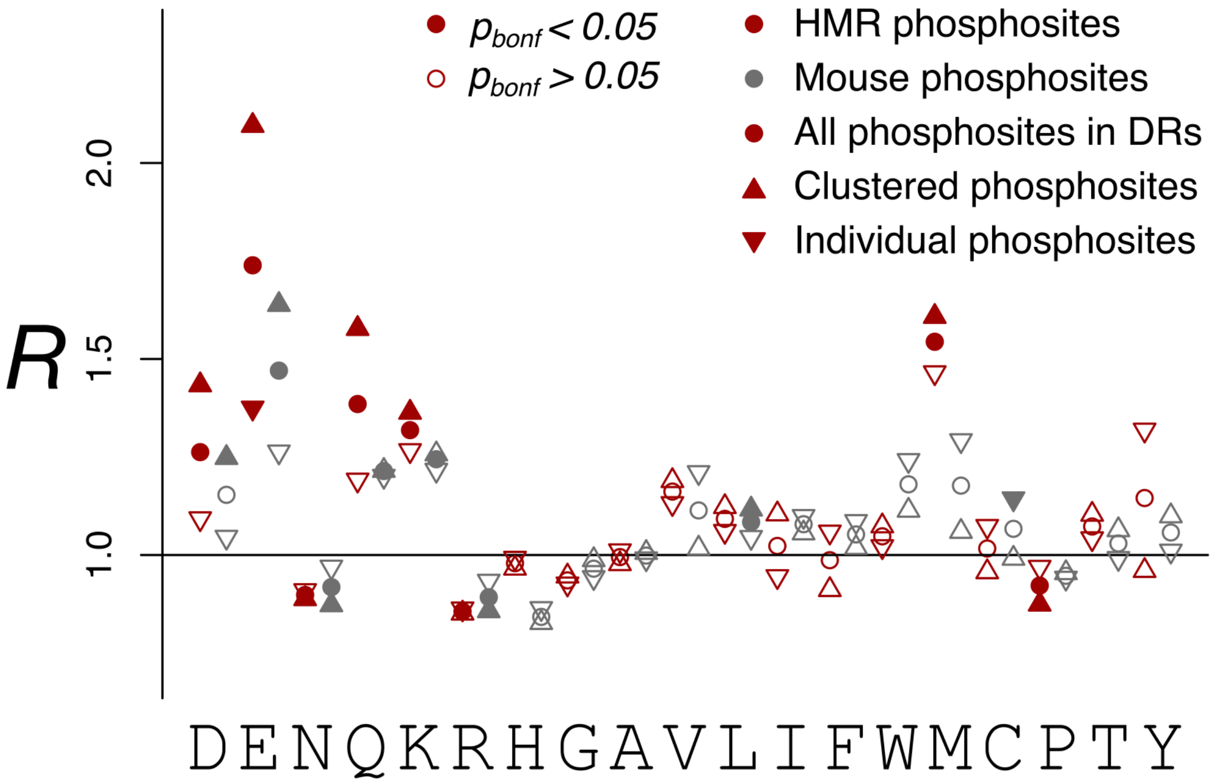


**Supplementary Figure S8 | *R* values of the pS-to-X mutations for different phosphosite sets.** Note the stronger effects observed for the HMR dataset compared to the mouse dataset and for clustered phosphoserines with respect to individual ones.

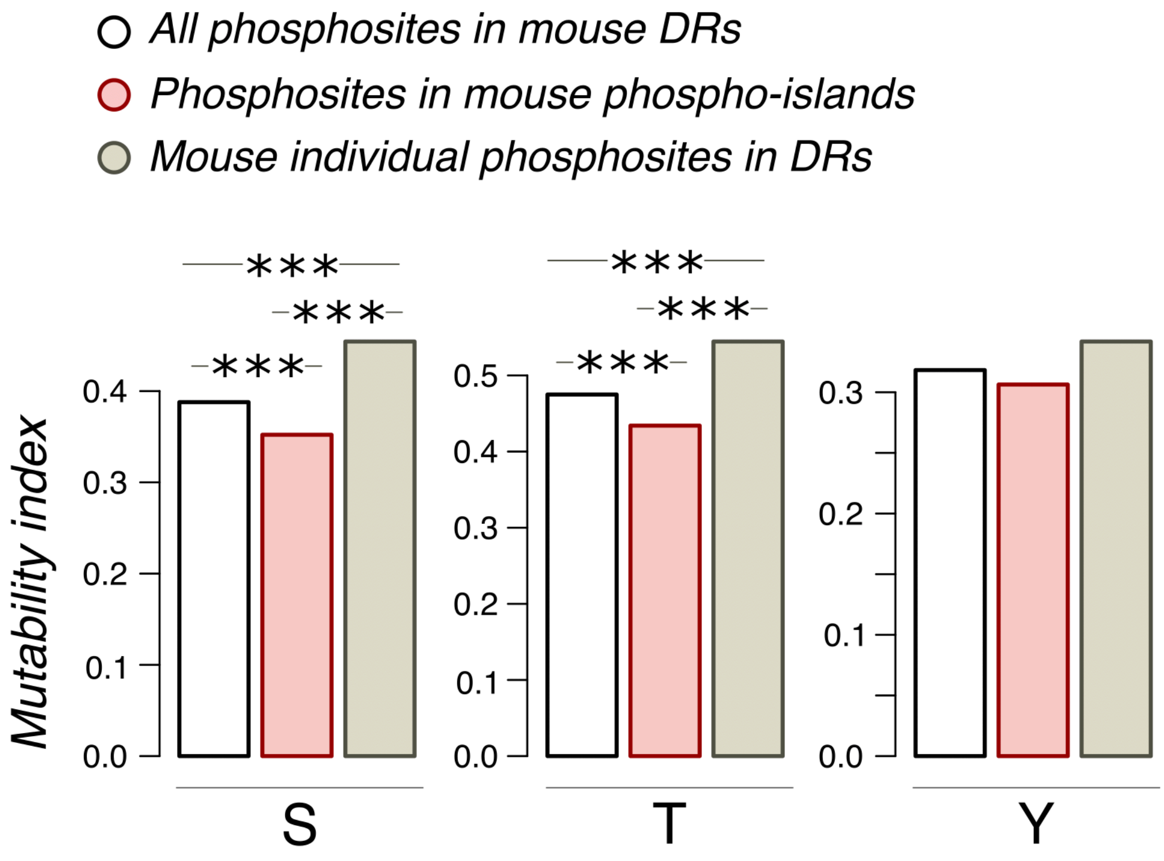

**Supplementary Figure S9 | Conservation of phosphorylated amino acids for various mouse datasets.** The mutability index is calculated for a phosphorylated amino acid with respect to its non-phosphorylated counterpart as the sum of probabilities of STY-to-X mutations calculated for all tree branches for the phosphorylated STY amino acid divided by the same sum for the respective non-phosphorylated amino acid. Three asterisks indicate statistically significant differences (*p* < 0.001, chi-squared contingency test).

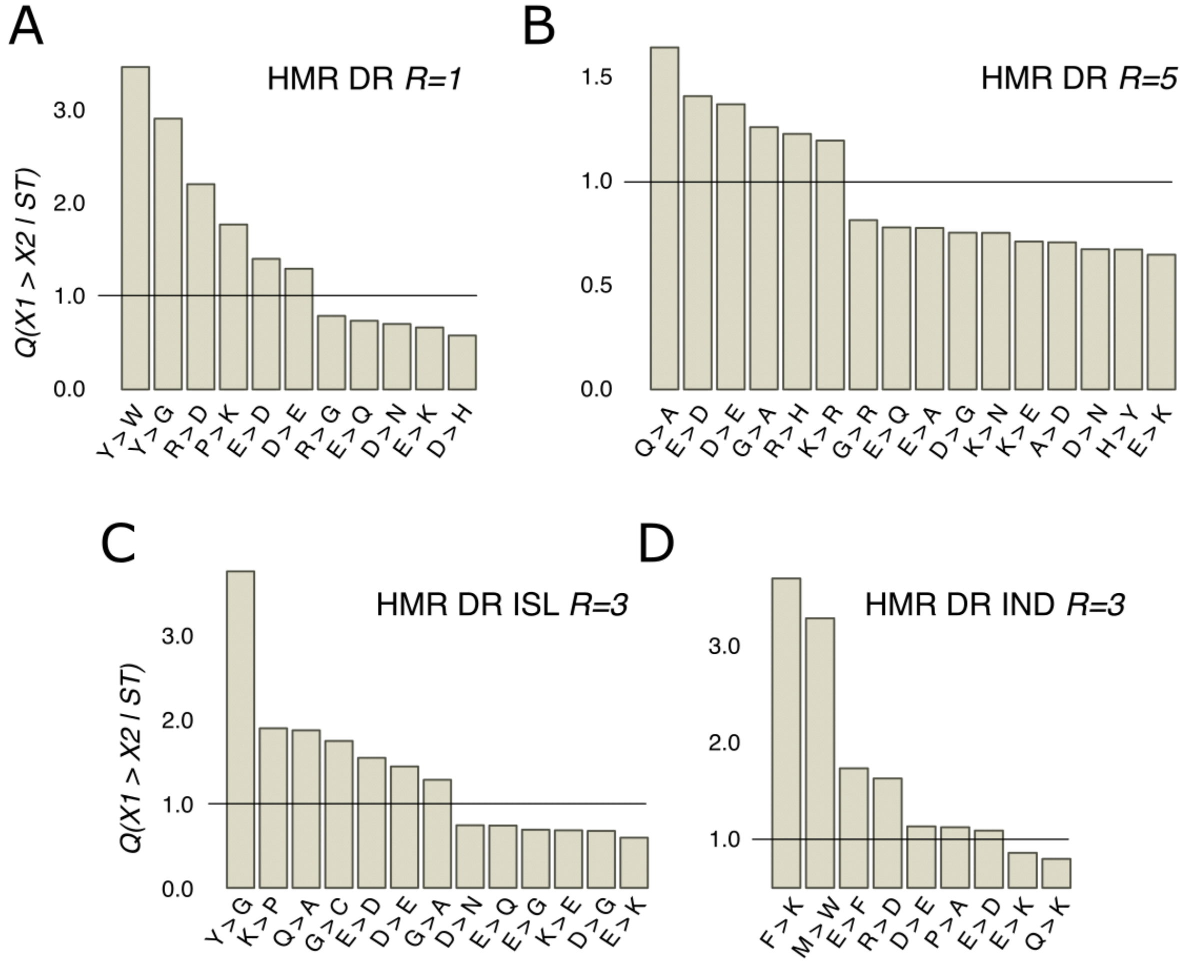


**Supplementary Figure S9 | *R* values of mutations near ST phosphosites with probabilities significantly different from the expected ones for various HMR subsets. (A)** HMR sites located in disordered regions with the window radius 1. **(B)** HMR sites located in disordered regions with the window radius 3. **(C)** Clustered HMR sites with the window radius 3. **(D)** Individual HMR sites with the window radius 3.

#### 2. Supplementary Tables

Supplementary tables are provided in the additional XLSX file.

**/hmr_sites_list**: List of all HMR-sites considered in the present study. Each site is marked as present in ordered/disordered region, having one of the five motif types, and being a part of a phosphosite cluster or individual. Phosphosites located in DRs are considered as being neither clustered nor individual.
**/mouse_site_list**: List of mouse dataset phosphosites.
**/hmr_psite_islands**: List of HMR phosphor-islands.

**/mouse_psite_islands**: List of mouse phosphor-islands.
**/R table**: Table with *R* values and the respective corrected *p*-values for all considered phosphosite datasets. Fig. 3 and Suppl. Figs. S2-S6 correspond to this table.
**/local_mutations**: Q values of contextual mutations for different HMR phosphosite datasets and different window radius. Corresponds to Fig. 5 and Suppl. Fig. S9.
**/phosphosite conservation**: Mutability statistics for the mouse dataset. Corresponds to Suppl. Fig. S8.
